# Patterns of Typical and Atypical Age-related Brainstem Volume losses

**DOI:** 10.64898/2026.05.21.726989

**Authors:** SG Mueller, RS Mackin

## Abstract

**Background:** The brainstem and its different sub-systems control essential functions such as motor agility etc. that worsen with age. The purpose of this study was: 1. To assess the impact of age-related volume loss within three brainstem sub-systems on functions supported by them. 2. To use data-driven machine learning to identify different volume loss patterns or subtypes and investigate how they are associated with function.

**Methods:** Structural MRI and behavioral data from 674 Human Connectome Project Aging (HCA) participants was used in this project. The brainstem was extracted, internal brainstem structures segmented and the segmentations warped onto a probabilistic population atlas on which the nuclei of interest had been labeled. Jacobian deformation maps were calculated, each roi’s mean Jacobians extracted and converted into z-scores with and without correction for age. Linear regression analyses were used to assess volume – function (cognition, motor agility, autonomic control) associations for each roi belonging to the sub-system supporting these functions. Subtype and Stage Inference (SuStaIn) was used to identify different volume loss patterns in each sub-system.

**Results:** Age explained larger percentage of the variation of the behavioral variables than brainstem volumes. SuStaIn identified up to 4 subtypes, one representing typical aging and the remainder atypical aging. The subtypes did not significantly differ behaviorally with the exception of grip strength and diastolic blood pressure.

**Conclusion:** Aging affects brainstem systems which contributes to the worsening of these functions with increasing age. SuStaIn detected different patterns of volume loss or subtypes within each of the brainstem systems.

## 1. Introduction

The brainstem consists of different subsystems that play an important role in the control of essential functions such as motor function, cognition, autonomic control etc.. These functions are known to be susceptible to the effects of age (Kocevska et al., 2021, Mander et al. 2017, Wan et al. 2023, Schumann et al., 2023, Baker et al., 2016, Harada et al., 2013, Austad et al., 2019, Lindenberger, 2014). But whereas numerous studies investigated how these functions are affected by age-related cortical and subcortical volume losses/atrophy (e.g., Gonzalez et al., 2024, Amorim et al.,2018, Holtzer et al., 2014, Rosano et al., 2007, Kong et al. 2023, De Looze et al. 2020), the number of studies trying to understand the contribution of age-related volume loss of brainstem structures in this process is relatively small. Furthermore, many of the studies that did investigate the role of the brainstem either assessed the brainstem as a whole or focused on its major divisions, i.e., mesencephalon, pons medulla oblongata, and did not attempt to distinguish between the different functional brainstem subsystems or nuclei within these major divisions (Hu et al., 2024, Dutt et al. 2020, Sander et al., 2019, Bouhrara et al., 2020). Finally, studies (Mueller 2023, Langley et al., 2022, van Egroo et al., 2022, Lambert et al., 2013, Xing et al. 2018) that identified individual nuclei usually investigated them in isolation and not as part of a system of brainstem structures supporting a specific function. The overall goal of this project was therefore: 1. To identify brainstem nuclei belonging to brainstem systems supporting specific functions such as cognition, motor function, in particular locomotion and strength, and autonomic control based on reports in the literature. 2. To segment and extract their volumes from the MRIs of a large population of normally aging individuals, establish the assumed structure/behavior relationship and investigate the impact of age on that relationship. 3. Use a recently developed unsupervised and data-driven machine learning algorithm called Subtype and Stage Inference or SuStaIn that employs a spatiotemporal clustering approach to disentangle subtypes with distinct volume patterns (Young et al. 2018, Aksman et al. 2021) to identify common or typical volume patterns as well as variants or atypical volume patterns within each system and to investigate how these variants influence the structure/behavior relationship.

## 2. Methods

### 2.1. Population

The imaging and behavioral data of 674 subjects (age mean(SD) : 58.8 (14.8), age range: 36 – 100, m/f: 292/382) from the Human Connectome Project Aging (HCA) (Bookheimer et al. 2019) was used for this project. The aim of HCA is to study “typical aging”, i.e., its population includes participants who exhibit health issues typically seen in their age cohort, e.g., hypertension, musculoskeletal pain, but do not suffer from pathological conditions, e.g., major depression, sleep apnea, stroke or mild cognitive impairment due to a neurodegenerative process etc.. It collects a variety of structural and functional MR images, as well as behavioral and biological data. The T1 and T2 images of these 674 subjects were used for this study as well as the following behavioral measures for those of that group for whom the data was available in the NDA database as of Sept 2022: cognition: Global cognitive function (total cognition composite score (with and without age correction, n = 508), list sorting working memory (with and without age correction n = 519), short term memory (Rey auditory verbal learning (rvlt) short delay total correct, raw scores and age adjusted residuals, n = 561). Motor function: Endurance (number of course turns during 2 min walk and normative standard score, n = 502), locomotion (time for 4 m walk, raw and computed score, n = 509), grip strength of dominant and non-dominant hand (grip) (with and without age correction, n = 517). Autonomic function: systolic and diastolic blood pressure while sitting (n = 570).

Studies using anonymized data where it is not possible to ascertain the subject’s identity are considered non-human subject studies and exempt from UCSF IRB review.

### 2.2. Imaging

All participants were scanned on a customized Siemens 3T “Connectome Skyra” at Washington University using a standard 32-channel Siemens receive and a body transmission coil (Bookheimer et al. 2019). The distortion corrected T1 weighted images (3D MPRAGE TR=2400 ms, TE=2.14 ms, TI=1000 ms, FA=8°, Bandwidth (BW)=210 Hz per pixel, Echo Spacing (ES) =7.6 ms, 0.8 mm isotropic resolution) and T2 weighted images (SPACE, TR=3200 ms, TE=565 ms, 0.8mm isotropic resolution with same matrix and slices as T1 images were used for this project.

### 2.3. Image processing

The details of the approach used for segmenting the internal brainstem structures are provided in Mueller, 2023. In short, k-means clustering was used to identify 5 probabilistic brainstem and diencephalon intensity clusters corresponding to 5 brainstem tissue types from a T1, T2 and T1/T2 ratio brainstem image. SPM12′s non-linear diffeomorphic mapping algorithm (DARTEL) was used to warp these segmentations onto a probabilistic brainstem tissue template in MNI space that had been generated from the brainstem segmentations from all 674 HCP subjects. The following brainstem regions of interest (roi) had been manually delineated on this template using the brainstem atlases from Naidich et al., 2009 and Paxinos and Huang, 2011 as references: periaqueductal gray (PAG), ventral tegmental area (VTA), raphe dorsalis ncl. (DR) median raphe ncl. (MedR), raphe magnus ncl. (MR), raphe obscurus (OR) and raphe pallidus ncl. (PR), left and right substantia nigra (SN), ncl ruber (NR), ncl. reticularis cuneiformis (CR), locus coeruleus (LC), ncl. pedunculopontinus (PP), ncl. reticularis pontis oralis (RPO), ncl. pontis caudalis (RPC), ncl. parabrachialis (PB), ncl. reticularis gigantocellularis and parvocellularis (RG), ncl. tractus solitarii (NTS), ncl. olivarius inferior (OI), and ventrolateral medulla (VLM). The transformation matrices generated during the warping step were converted into ICV corrected Jacobian determinant maps from which the mean intensities from each roi were extracted for each subject. Based on reports in the literature, the following rois were identified as significantly contributing to either cognition, motor or autonomic control:

1. Cognitive control system of the brainstem or cognitive brainstem: PAG, VTA, LC, DR, MedR (Benarroch,2012, Benarroch, 2023, Ramirez et al.,2023, Zhang et al. 2018)
2. Motor control brainstem system or motor brainstem: SN, NR, PP, CR. RPO, RPC, OI, RG (Capelli et al. 2017, Leiras et al. 2022, Brownstone et al. 2018, Boudry et al., 2014, Noga et al., 2022, Ali and Benarroch, 2022)
3. Autonomic control/blood pressure control brainstem system or autonomic brainstem: PAG, PB, NTS, VLM, MR, OR, PR (Benarroch, 2019, Benarroch, 2020, Guyenet et al., 2022, Braun et al., 2024, Kishi et al., 2023)

Each system roi’s mean Jacobian was extracted and converted into z-scores with (residuals obtained from linear regression with age, input used for subtyping and staging) and without age correction (z-scores calculated using mean and standard deviation from all 674 subjects, input used to investigate relationships with age and behavior).

### 2.4. Subtype and Stage Inference (SuStaIn)

Three SuStaIn analyses, i.e., one for each system, were performed. For each of them, the mean Jacobians from each roi in the system were extracted and age corrected z-scores (defined so that higher values indicated more pronounced volume loss) calculated for all 674 subjects. The piecewise linear z-score model of the SuStaIn algorithm (Young et al., 2018, Aksman et al.,2021) with 25 start points, 1,000,000 Markov Chain Monte Carlo iterations and evaluating up to 4 clusters or subtypes was ran to identify distinct subtypes and distinct stages within each of these subtypes. This was followed by 10-fold cross-validation to evaluate the optimal number of subtypes (defined as cluster number with lowest cross-validation information criterion (CVIC)) that best describe unseen data and to assess the stability of subtypes across folds. Each of the SuStaIn subtypes and each of the stages is expressed to some degree (indicated by a maximum likelihood value ranging from 0-1) in each participant. Using a maximum likelihood cut-off >0.5 for models with 2 and >0.33 for models with 3 subtypes and identifying the stage with the highest maximum likelihood within this subtype, each participant’s age corrected brainstem z-scores were assigned to a subtype and a stage. Participants without volume abnormalities in the system of interest were assigned to subtype 0, stage 0, reflecting the normal or typical aging process in this system. Participants with brainstem volume loss (z-scores > 1) within the system of interest were assigned to one of the atypical aging subtypes with stages 1 and higher.

### 2.5. Statistics

Linear regression (with and without age co-variate) were used to assess associations between behavior (cognition, motor, blood pressure) and roi volumes. Holm’s correction was used to correct for multiple comparisons. One-way ANOVA (Welch’s test) followed by Games Howell post hoc tests were used to assess differences of behavioral variables (age corrected version) between subtypes.

### 2.6. Research Data

The imaging and behavioral data used in this project came from the publicly available Human Connectome Project of Aging. The Matlab scripts and priors used for this project will be made available to qualified researchers upon request.

## 3. Results

Please see Table 1 for a summary of the study population’s behavioral characteristics.

**Table 1.** Behavioral Variables: Population Overview.

|  | **N** | **mean (SD)** |
| --- | --- | --- |
| Total cognition composite (no age) | 519 | 106 (11.3) |
| Total cognition composite (age adjust) | 508 | 109 (15.7) |
| Working memory (no age) | 519 | 101 (11.2) |
| Working memory (age adjust) | 519 | 100 (23.6) |
| Rvlt short delay total correct (no age) | 561 | 61.3 (14.1) |
| Rvlt short delay total correct (age adjust) | 561 | 0.000 (13.2) |
| Endurance (no age) | 502 | 11.9 (1.9) |
| Endurance (age adjust) | 502 | 101 (12.7) |
| Locomotion (no age) | 508 | 3.3 (0.6) |
| Locomotion (age adjust) | 508 | 1.3 (0.2) |
| Grip strength dominant (no age) | 517 | 72.7 (23.6) |
| Grip strength dominant (age adjust) | 517 | 100 (20) |
| Grip strength non-dominant (no age) | 517 | 69.4 (23.3) |
| Grip strength non-dominant (age adjust) | 517 | 99.5 (20.3) |
| Systolic blood pressure | 566 | 128 (15.6) |
| Diastolic blood pressure | 566 | 78.8 (10.4) |

Rvlt, Rey auditory verbal learning short delay total correct

In 216 HCP subjects (mean age (SD) : 59.1 (14.1), range 36.4 – 100 years of age) the volume losses were within the normal aging range (subtype 0) within all three systems. 145 HCP subjects (mean age (SD) : 59.1 (15.2), range 36.4 – 100 years of age) had volume losses exceeding normal aging in 1 system, 184 (mean age (SD) : 58.1 (14.8), range 36.0 – 89.1 years of age) had volume losses exceeding normal aging in 2 systems, and 129 (mean age (SD) : 58.1 (15.8) range 36.3 – 100 years of age) had volume losses exceeding normal aging in 3 systems.

### 3.1.1. Cognitive brainstem system

Please see Table 2. Total cognition composite score, list sorting working memory short term memory performance and Rey auditory short delay total correct scores worsened with increasing age. Age alone explained 8 – 12.6% of the variation of these variables. Strong, i.e., surviving correction for multiple comparisons, volume – cognition associations without correction for age were found for VTA and LC. VTA and LC volumes explained between 1 – 15.7% of the variation of these variables. When age was also included into the model, only VTA associations survived corrections for multiple comparisons.

**Table 2.** Relationship Cognitive Function with brainstem structures involved in cognition.

|  |  |  | Total cognition<br>composite | List sorting<br>working memory | Rvlt short delay<br>total correct |
| --- | --- | --- | --- | --- | --- |
| Age | Estimate |  | **-0.225** | **-0.245** | **-0.355** |
|  | Rsquare |  | **0.076** | **0.094** | **0.126** |
|  | p |  | **<0.001** | **<0.001** | **<0.001** |
| PAG | Estimate |  | -0.274 | 0.311 | 1.030 |
|  | Rsquare |  | 0.001 | 0.001 | 0.005 |
|  | p |  | 0.058 | 0.527 | 0.081 |
| VTA | Estimate |  | **-1.420** | **-1.390** | **-3.070** |
|  | Rsquare |  | **0.016** | **0.157** | **0.048** |
|  | p |  | **0.004** | **0.004** | **<0.001** |
| LC left | Estimate |  | -0.771 | 0.274 | **-1.650** |
|  | Rsquare |  | 0.004 | 0.000 | **0.013** |
|  | p |  | 0.132 | 0.797 | **0.006** |
| LC right | Estimate |  | *-1.130* | **-1.530** | **-1.930** |
|  | Rsquare |  | *0.010* | **0.018** | **0.018** |
|  | p |  | *0.025* | **0.002** | **0.001** |
| MedR | Estimate |  | -0.320 | -0.137 | 0.154 |
|  | Rsquare |  | 0.001 | 0.000 | 0.000 |
|  | p |  | 0.527 | 0.783 | 0.797 |
| DR | Estimate |  | 0.551 | 0.318 | -0.273 |
|  | Rsquare |  | 0.002 | 0.001 | 0.000 |
|  | p |  | 0.265 | 0.515 | 0.643 |
| PAG | Estimate | vol | -0.930 | -0.377 | 0.134 |
|  | Rsquare adjusted | vol&age | 0.081 | 0.092 | 0.123 |
|  | p | vol | 0.057 | 0.431 | 0.812 |
| VTA | Estimate | vol | -0.572 | -0.453 | **-1.733** |
|  | Rsquare adjusted | vol&age | 0.077 | 0.092 | **0.137** |
|  | p | vol | 0.247 | 0.349 | **0.003** |
| LC left | Estimate | vol | -0.029 | -0.176 | -0.534 |
|  | Rsquare adjusted | vol&age | 0.744 | 0.091 | 0.125 |
|  | p | vol | 0.955 | 0.723 | 0.358 |
| LC right | Estimate | vol | -0.303 | -0.655 | -0.656 |
|  | Rsquare adjusted | vol&age | 0.037 | 0.094 | 0.125 |
|  | p | vol | 0.548 | 0.184 | 0.265 |
| MedR | Estimate | vol | 0.382 | 0.635 | *1.132* |
|  | Rsquare adjusted | vol&age | 0.075 | 0.093 | *0.129* |
|  | p | vol | 0.442 | 0.192 | *0.047* |
| DR | Estimate | vol | 0.713 | 0.492 | -0.073 |
|  | Rsquare adjusted | vol&age | 0.078 | 0.093 | 0.123 |
|  | p | vol | 0.134 | 0.290 | 0.895 |

bold, corrected for multiple comparisons with Holm's test, italics, significant without correction for multiple comparisons PAG, periaqueductal gray; VTA, ventral tegmental area; LC, locus coeruleus; MedR, median raphe; DR, dorsal raphe rvlt, rey auditory verbal learning

### 3.1.2. SuStaIn

Please see Figure 1 and supplementary Table 1. SuStaIn identified two subtypes with increasing volume loss indicating atypical aging. In subtype 1 mild volume loss (z-scores 1-1.99) started in the MedR followed by DR and PAG. VTA and LC volume loss occurred only in later stages when the volume loss in the other regions was severe (z-scores>2). Subtype 2 was characterized by early mild volume loss in LC and VTA where it progressed to severe volume losses in the late stages without spreading to other regions.

**Figure 1.**
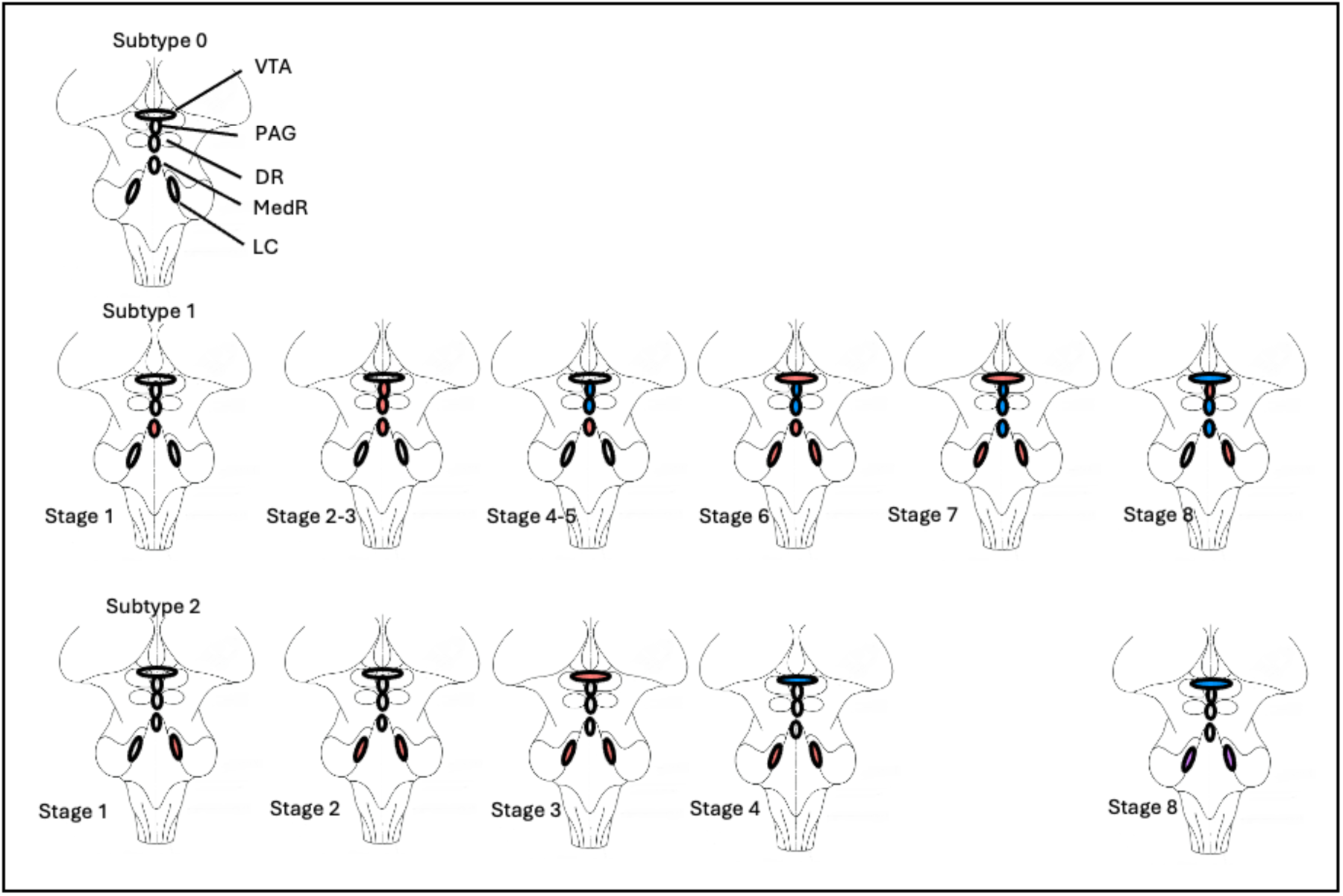
Subtypes and subtype stages of the cognitive control system. Subtype 0 represents normal aging. Subtypes 1 and 2 represent atypical aging. VTA, ventral tegmental area, PAG, periaqueductal gray, LC, locus coeruleus, DR, nucleus raphe dorsalis, MedR, nucleus raphe medianus. Red indicates volume losses, i.e., age corrected z-scores >1 but < 2, blue indicates volume losses with age corrected z-scores ≥ 2 but < 3, violet indicates volume losses with age corrected z-scores ≥ 3. Please see main text for a description of the pattern.

Age adjusted total cognition composite scores, list sorting working memory and Rey auditory short delay total correct scores were more impaired in subtype 2 but these differences did not reach significance, see Table 3.

**Table 3.** Cognition by Subtypes.

|  | Subtype | N | Mean | SD | ANOVA p-value |
| --- | --- | --- | --- | --- | --- |
| Age | 0 | 405 | 59.14 | 14.62 | 0.733 |
|  | 1 | 132 | 58.20 | 14.55 |  |
|  | 2 | 137 | 58.23 | 15.81 |  |
| Total cognition<br>composite score | 0 | 305 | 109.42 | 15.73 | 0.425 |
|  | 1 | 99 | 109.76 | 14.84 |  |
|  | 2 | 104 | 107.19 | 16.41 |  |
| List sorting working<br>memory | 0 | 314 | 101.34 | 22.07 | 0.391 |
|  | 1 | 99 | 98.69 | 19.64 |  |
|  | 2 | 106 | 98.18 | 30.22 |  |
| Rvlt short delay<br>total correct | 0 | 333 | 0.76 | 13.36 | 0.141 |
|  | 1 | 115 | -0.23 | 12.98 |  |
|  | 2 | 113 | -2.01 | 12.63 |  |

### 3.2.1. Motor control brainstem system

Please see Table 4. Endurance, speed and grip strength worsened with increasing age. Age explained between 4-18.5% of the variability of these parameters. Without correcting for age strong associations with endurance, speed and strength indicating a loss of function with volume loss were found for SN, NR, CR, RPO and OI. Volume loss explained between 2 – 7% of the variability of endurance, speed etc. RG volumes were strongly associated with hand strength but not in the expected direction, i.e., grip strength increased with more pronounced volume loss. Correcting for age weakened the associations and only those for SN, RG and RPO survived correction for multiple comparisons. RG and newly also CR and RPO volumes were positively associated with hand grip strength, i.e., even after correction for age effects grip strength increased with volume loss.

**Table 4.** Relationship Motor function with brainstem structures involved in motor cntrol.

|  |  | Endurance<br>No course<br>turns during<br>2 min walk | Locomotion<br>time for 4<br>meter walk | Grip<br>Strength<br>dominant<br>hand | Grip Strength<br>non-dominant<br>hand |
| --- | --- | --- | --- | --- | --- |
| Age | Estimate | **-0.057** | **0.008** | **-0.409** | **-0.367** |
|  | Rsquare | **0.185** | **0.036** | **0.059** | **0.049** |
|  | p | **<0.001** | **<0.001** | **<0.001** | **<0.001** |
| SN left | Estimate | **-0.502** | **0.082** | -1.460 | -105.0 |
|  | Rsquare | **0.073** | **0.019** | 0.004 | 0.002 |
|  | p | **<0.001** | **0.002** | 0.164 | 0.310 |
| SN right | Estimate | **-0.413** | **0.090** | -0.920 | -0.609 |
|  | Rsquare | **0.048** | **0.021** | 0.001 | 0.001 |
|  | p | **<0.001** | **0.001** | 0.395 | 0.570 |
| NR left | Estimate | **-0.428** | **0.090** | -1.130 | -0.713 |
|  | Rsquare | **0.0529** | **0.023** | 0.002 | 0.001 |
|  | p | **<0.001** | **<0.001** | 0.276 | 0.485 |
| NR right | Estimate | **-0.483** | **0.089** | -0.980 | -0.663 |
|  | Rsquare | **0.064** | **0.022** | 0.002 | 0.001 |
|  | p | **<0.001** | **<0.001** | 0.349 | 0.522 |
| PP left | Estimate | -0.096 | 0.026 | -0.966 | -0.848 |
|  | Rsquare | 0.003 | 0.002 | 0.002 | 0.001 |
|  | p | 0.255 | 0.332 | 0.359 | 0.416 |
| PP right | Estimate | 0.001 | -0.004 | -0.069 | -0.282 |
|  | Rsquare | 0.000 | 0.000 | 0.000 | 0.000 |
|  | p | 0.929 | 0.894 | 0.948 | 0.788 |
| CR left | Estimate | **-0.241** | 0.029 | 0.526 | 0.793 |
|  | Rsquare | **0.014** | 0.002 | 0.001 | 0.001 |
|  | p | **0.004** | 0.287 | 0.621 | 0.451 |
| CR right | Estimate | **-0.331** | 0.041 | -0.252 | 0.163 |
|  | Rsquare | **0.030** | 0.004 | 0.000 | 0.000 |
|  | p | **<0.001** | 0.128 | 0.813 | 0.877 |
| RPO left | Estimate | **-0.018** | 0.067 | 1.680 | 1.600 |
|  | Rsquare | **-0.292** | 0.126 | 0.005 | 0.005 |
|  | p | **<0.001** | 0.011 | 0.105 | 0.120 |
| RPO right | Estimate | **-0.275** | 0.058 | 0.976 | 0.694 |
|  | Rsquare | **0.022** | 0.010 | 0.002 | 0.001 |
|  | p | **<0.001** | 0.028 | 0.345 | 0.497 |
| RPC left | Estimate | -0.138 | -0.003 | -0.023 | -0.051 |
|  | Rsquare | 0.005 | 0.000 | 0.000 | 0.000 |
|  | p | 0.103 | 0.908 | 0.983 | 0.961 |
| PRC right | Estimate | -0.161 | 0.013 | 1.260 | 1.190 |
|  | Rsquare | 0.007 | 0.000 | 0.003 | 0.003 |
|  | p | 0.058 | 0.627 | 0.233 | 0.255 |
| RG left | Estimate | 0.082 | 0.016 | **5.780** | **5.640** |
|  | Rsquare | 0.002 | 0.001 | **0.061** | **0.059** |
|  | p | 0.321 | 0.554 | **<0.001** | **<0.001** |
| RG right | Estimate | 0.070 | 0.029 | **5.520** | **5.220** |
|  | Rsquare | 0.001 | 0.002 | **0.054** | **0.049** |
|  | p | 0.400 | 0.273 | **<0.001** | **<0.001** |
| OI left | Estimate | **-0.339** | 0.016 | -2.290 | -1.560 |
|  | Rsquare | **0.035** | 0.001 | 0.010 | 0.005 |
|  | p | **<0.001** | 0.527 | 0.025 | 0.122 |
| OI right | Estimate | **-0.324** | 0.004 | -2.550 | -1.750 |
|  | Rsquare | **0.031** | 0.000 | 0.012 | 0.005 |
|  | p | **<0.001** | 0.868 | 0.014 | 0.088 |
| SN left | Estimate vol | **-0.247** | **-0.250** | -0.024 | 0.712 |
|  | Rsquare adj vol & age | **0.197** | **0.197** | 0.034 | 0.056 |
|  | p vol | **0.002** | **<0.001** | 0.034 | 0.513 |
| SN right | Estimate vol | -0.131 | 0.053 | 1.615 | 1.706 |
|  | Rsquare adj vol & age | 0.185 | 0.038 | 0.059 | 0.049 |
|  | p vol | 0.121 | 0.073 | 0.155 | 0.131 |
| RN left | Estimate vol | -0.139 | 0.054 | 1.401 | -0.414 |
|  | Rsquare adj vol & age | 0.179 | 0.039 | 0.0587 | 0.049 |
|  | p vol | 0.091 | 0.056 | 0.201 | 0.138 |
| RN right | Estimate vol | -0.176 | 0.049 | 1.887 | 1.964 |
|  | Rsquare adj vol & age | **0.181** | 0.037 | 0.061 | 0.051 |
|  | p vol | **0.038** | 0.088 | 0.094 | 0.080 |
| PP left | Estimate vol | 0.0151 | 0.011 | -0.263 | -0.216 |
|  | Rsquare adj vol & age | 0.181 | 0.032 | 0.056 | 0.045 |
|  | p vol | 0.844 | 0.682 | 0.798 | 0.883 |
| PP right | Estimate vol | 0.107 | -0.016 | 1.033 | 0.238 |
|  | Rsquare adj vol & age | 0.185 | 0.032 | 0.056 | 0.045 |
|  | p vol | 0.165 | 0.548 | 0.621 | 0.817 |
| CR left | Estimate vol | 0.085 | -0.018 | **3.227** | **3.274** |
|  | Rsquare adj vol & age | 0.183 | 0.033 | **0.071** | **0.061** |
|  | p vol | 0.309 | 0.523 | **0.004** | **0.003** |
| CR right | Estimate vol | 0.011 | -0.009 | 2.672 | 2.872 |
|  | Rsquare adj vol & age | 0.181 | 0.032 | 0.066 | 0.057 |
|  | p vol | 0.895 | 0.768 | 0.018 | 0.011 |
| RPO left | Estimate vol | -0.059 | 0.361 | **3.782** | **3.491** |
|  | Rsquare adj vol & age | 0.182 | 0.035 | **0.0792** | **0.066** |
|  | p vol | 0.455 | 0.183 | **<0.001** | **<0.001** |
| RPO right | Estimate vol | -0.014 | 0.023 | **3.277** | 2.737 |
|  | Rsquare adj vol & age | 0.181 | 0.033 | **0.0731** | 0.058 |
|  | p vol | 0.861 | 0.408 | **0.002** | 0.010 |
| RPC left | Estimate vol | -0.006 | -0.022 | 0.977 | 0.847 |
|  | Rsquare adj vol & age | 0.181 | 0.033 | 0.057 | 0.047 |
|  | p vol | 0.939 | 0.407 | 0.348 | 0.414 |
| PRC right | Estimate vol | -0.023 | -0.006 | 2.308 | 2.134 |
|  | Rsquare adj vol & age | 0.182 | 0.032 | 0.065 | 0.053 |
|  | p vol | 0.771 | 0.822 | 0.27 | 0.040 |
| RG left | Estimate vol | 0.102 | 0.013 | **6.005** | **5.841** |
|  | Rsquare adj vol & age | 0.184 | 0.032 | **0.121** | **0.109** |
|  | p vol | 0.173 | 0.616 | **<0.001** | **<0.001** |
| RG right | Estimate vol | 0.087 | 0.027 | **5.683** | **5.365** |
|  | Rsquare adj vol & age | 0.184 | 0.034 | **0.113** | **0.098** |
|  | p vol | 0.250 | 0.291 | **<0.001** | **<0.001** |
| OI left | Estimate vol | -0.024 | -0.034 | 0.0004 | 0.596 |
|  | Rsquare adj vol & age | 0.182 | 0.035 | 0.056 | 0.046 |
|  | p vol | 0.767 | 0.224 | 0.997 | 0.580 |
| OI right | Estimate vol | 0.013 | -0.052 | -0.185 | 0.494 |
|  | Rsquare adj vol & age | 0.181 | 0.038 | 0.056 | 0.046 |
|  | p vol | 0.880 | 0.069 | 0.867 | 0.654 |

bold, corrected for multiple comparisons with Holm's test, italics, significant without correction for multiple comparisons SN, substantia nigra; PP, ncl pedunculo pontinus; RG, ncl. reticularis gigante- and parvocellularis; CR, ncl. cuneiformis; RPO, ncl reticularis pontis oralis, RPC, ncl pontis caudalis, NR, ncl ruber, OI ncl olivarius inferior

### 3.2.2. SuStaIn

Please see Figure 2 and supplementary Table 2. Three subtypes showed progressive volume loss indicating atypical aging. Subtype 1 was characterized by mild volume loss in OI and PP in the early stages that spread to CR, RPC and SN in later stages. Subtype 2 started with mild volume loss in RG that spread to RPC, RPO, CR and to SN in the later stages. Subtype 3 started in NR followed by SN and RPO. The volume loss became severe in SN in the later stages. Increased dominant (ANOVA p = 0.019) and non-dominant grip (ANOVA p = 0.007) strengths of subjects assigned to subtype 2 compared to subtype 0 were the only behavioral differences reaching significance (dominant: subtype 0: 97.67 (20.40) vs. subtype 2: 104.51 (17.3), p = 0.009; non-dominant: subtype 0: 96.97 (21.09) vs. subtype 2: 104.65 (17.71), p = 0.003). Please see Table 5.

**Figure 2.**
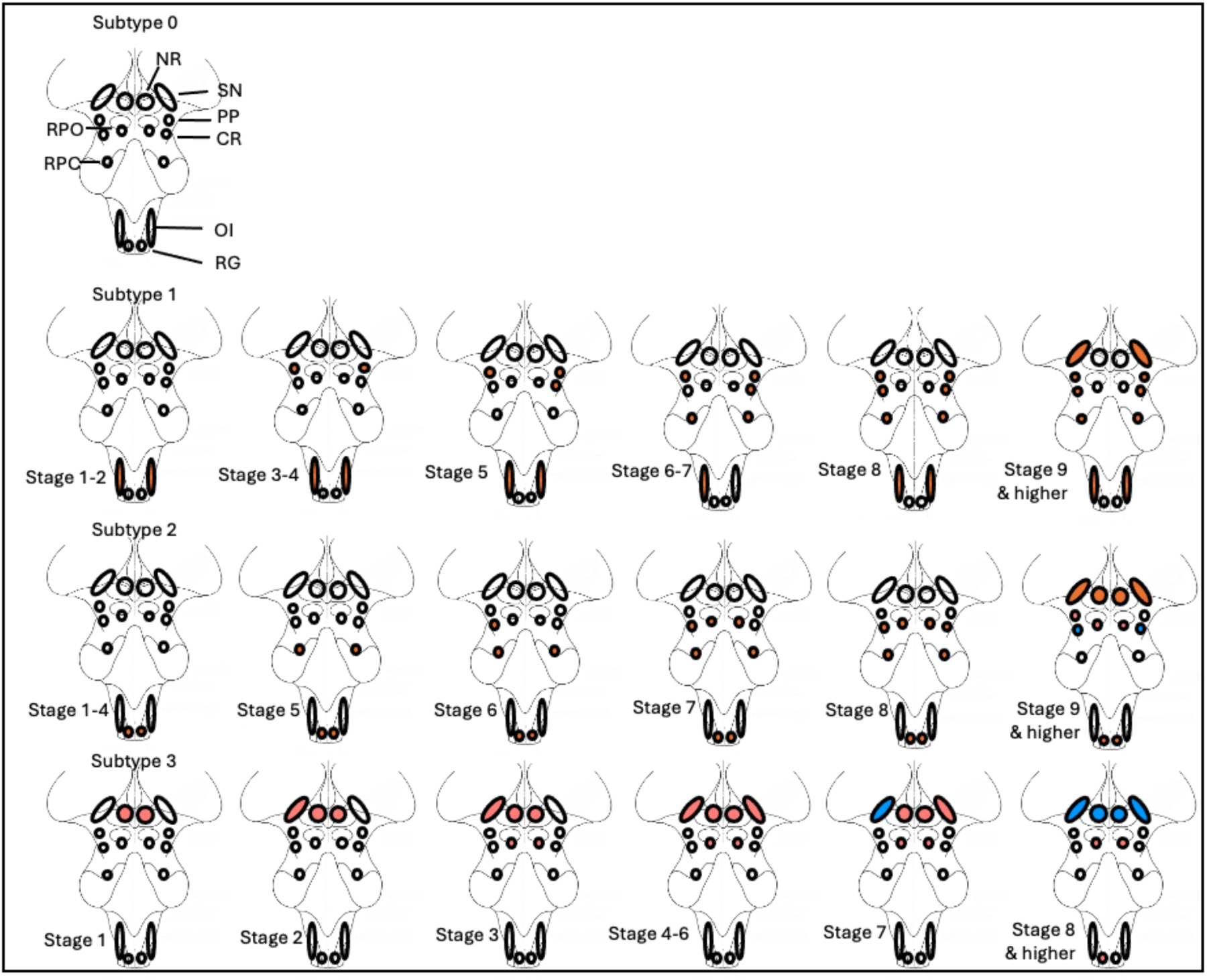
Subtypes and subtype stages of the motor control system. Subtype 0 represents normal aging. Subtypes 1, 2 and 3 represent atypical aging. NR, nucleus ruber, SN, substantia nigra, PP nucleus pedunculopontinus, CR, nucleus cuneiformis, RPO nucleus reticularis pontis oralis, RPC, nucleus reticularis pontis caudalis, OI, nucleus olivarius inferior, RG, nucleus gigantocellularis. Red indicates volume losses, i.e., age corrected z-scores >1 but < 2, blue indicates volume losses with age corrected z-scores ≥ 2 but < 3, violet indicates volume losses with age corrected z-scores ≥ 3. Please see main text for a description of the pattern.

**Table 5.** Motor Function by Subtypes.

|  | Subtype | N | Mean | SD | ANOVA p-value |
| --- | --- | --- | --- | --- | --- |
| age | 0 | 336 | 59.10 | 14.19 | 0.331 |
|  | 1 | 133 | 58.45 | 14.66 |  |
|  | 2 | 135 | 59.89 | 15.70 |  |
|  | 3 | 70 | 55.66 | 16.38 |  |
| Endurance -Normative Standard Score | 0 | 251 | 100.54 | 12.04 | 0.064 |
|  | 1 | 101 | 99.98 | 13.86 |  |
|  | 2 | 95 | 103.02 | 13.26 |  |
|  | 3 | 55 | **97.31** | 12.03 |  |
| Locomotion - 4 meter walk - Computed Score | 0 | 253 | 1.27 | 0.22 | 0.129 |
|  | 1 | 103 | **1.30** | 0.26 |  |
|  | 2 | 97 | 1.26 | 0.21 |  |
|  | 3 | 55 | 1.19 | 0.29 |  |
| Age-Corrected Standard Scores Dominant | 0 | 258 | 97.67 | 20.40 | 0.019 |
|  | 1 | 105 | 100.96 | 20.17 |  |
|  | 2 | 99 | **104.51** | 17.35 |  |
|  | 3 | 55 | 101.05 | 21.16 |  |
| Age-Corrected Standard Scores Non-Dominant | 0 | 258 | 96.97 | 21.09 | 0.007 |
|  | 1 | 105 | 101.19 | 19.72 |  |
|  | 2 | 99 | **104.65** | 17.71 |  |
|  | 3 | 55 | 99.31 | 20.39 |  |

**Table 6.** Associations of brainstem structures involved in autonomic control with blood pressure.

|  |  |  | systolic (sitting) | diastolic (sitting) |
| --- | --- | --- | --- | --- |
| Age | Estimate |  | **0.352** | -0.036 |
|  | Rsquare |  | **0.101** | 0.002 |
|  | p |  | **<0.001** | 0.248 |
| PAG | Estimate |  | 0.985 | -0.481 |
|  | Rsquare |  | 0.004 | 0.002 |
|  | p |  | 0.134 | 0.270 |
| PB left | Estimate |  | -0.096 | -0.429 |
|  | Rsquare |  | 0.000 | 0.002 |
|  | p |  | 0.884 | 0.328 |
| PB right | Estimate |  | -1.090 | -0.906 |
|  | Rsquare |  | 0.005 | *0.008* |
|  | p |  | 0.099 | *0.039* |
| NTS left | Estimate |  | **2.680** | -0.264 |
|  | Rsquare |  | **0.030** | 0.001 |
|  | p |  | **<0.001** | 0.545 |
| NTS right | Estimate |  | **2.610** | -0.349 |
|  | Rsquare |  | **0.028** | 0.001 |
|  | p |  | **<0.001** | 0.422 |
| VLM left | Estimate |  | *1.320* | *0.986* |
|  | Rsquare |  | *0.007* | *0.009* |
|  | p |  | *0.043* | *0.023* |
| VLM right | Estimate |  | *1.550* | *0.996* |
|  | Rsquare |  | *0.010* | *0.010* |
|  | p |  | *0.017* | *0.020* |
| MR | Estimate |  | 0.579 | -1.020 |
|  | Rsquare |  | 0.001 | *0.010* |
|  | p |  | 0.380 | *0.020* |
| OR | Estimate |  | *1.390* | -0.415 |
|  | Rsquare |  | *0.008* | 0.002 |
|  | p |  | *0.033* | 0.337 |
| PR | Estimate |  | 0.353 | **1.000** |
|  | Rsquare |  | 0.001 | **0.013** |
|  | p |  | 0.523 | **0.006** |
| PAG | Estimate | vol | 0.105 | -0.395 |
|  | Rsquare adjusted | vol&age | 0.098 | 0.001 |
|  | p | vol | 0.869 | 0.363 |
| PB left | Estimate | vol | -0.179 | -0.427 |
|  | Rsquare adjusted | vol&age | 0.098 | 0.002 |
|  | p | vol | 0.776 | 0.331 |
| PB right | Estimate | vol | -0.984 | -0.917 |
|  | Rsquare adjusted | vol&age | 0.102 | *0.008* |
|  | p | vol | 0.177 | *0.036* |
| NTS left | Estimate | vol | 0.816 | -0.054 |
|  | Rsquare adjusted | vol&age | 0.103 | 0.000 |
|  | p | vol | 0.231 | 0.899 |
| NTS right | Estimate | vol | 0.772 | -0.172 |
|  | Rsquare adjusted | vol&age | 0.100 | 0.000 |
|  | p | vol | 0.255 | 0.692 |
| VLM left | Estimate | vol | 0.636 | *1.080* |
|  | Rsquare adjusted | vol&age | 0.100 | *0.011* |
|  | p | vol | 0.312 | *0.012* |
| VLM right | Estimate | vol | 0.614 | *1.110* |
|  | Rsquare adjusted | vol&age | 0.100 | *0.010* |
|  | p | vol | 0.328 | *0.010* |
| MR | Estimate | vol | -0.287 | -0.978 |
|  | Rsquare adjusted | vol&age | 0.100 | *0.008* |
|  | p | vol | 0.651 | *0.025* |
| OR | Estimate | vol | 0.564 | -0.343 |
|  | Rsquare adjusted | vol&age | 0.100 | 0.002 |
|  | p | vol | 0.368 | 0.427 |
| PR | Estimate | vol | 0.347 | **1.210** |
|  | Rsquare adjusted | vol&age | 0.098 | **0.014** |
|  | p | vol | 0.508 | **0.005** |

bold, corrected for multiple comparisons with Holm's test, italics, significant without correction for multiple comparisons
PAG, periaqueductal gray, PB, parabrachial nucleus, NTS nucleus tractus solitarius, VLM, ventrolateral medulla, MR, raphe magna nucleus, OR, raphe obscurus nucleus, PR, raphe pallidus nucleus

### 3.3.1. Autonomic control/blood pressure control system

Please see Table 5. Systolic blood pressure increased with age. The blood pressure of the 566 subjects for whom that information was available was elevated (systolic 128 (15.6) mm Hg, diastolic 78.8 (10.4) mmHg diastolic) but not within the hypertensive range according the 2017 guidelines of the American Heart Association. The systolic (135.4 (17.5) mmHg) but not the diastolic (79.5 (12.5) mmHg) blood pressure of the subset of patients (n= 66) receiving a blood pressure lowering medication was within the hypertensive range and significantly higher (p = <0.001) than that of the participants not taking blood pressure lowering medication (126.6 (15.1) resp 78.7 (10.1) mmHg). Age explained about 10% of the variation of the systolic blood pressure. Diastolic blood pressure was not associated with age.

Volume loss within brainstem structures involved in autonomic control explained between 1- 3% of the blood pressure variation. Strong correlations with systolic blood pressure were found for NTS but got weaker after correcting for age. PR volumes were strongly correlated with diastolic blood pressure.

### 3.3.2. SuStaIn

Please see Figure 3 and supplementary Table 3. SuStaIn identified two subtypes with progressive volume loss. Subtype 1 was characterized by mostly mild volume loss in RO and VLM in the early stages that became severe and spread to NTS and RP in the only subject assigned to a late stage. In subtype 2 mild atrophy appeared in NTS in stage 2 but spread to MR, RP, PB and PAG in stages 2 and 4 and became severe in stage 8.

**Figure 3.**
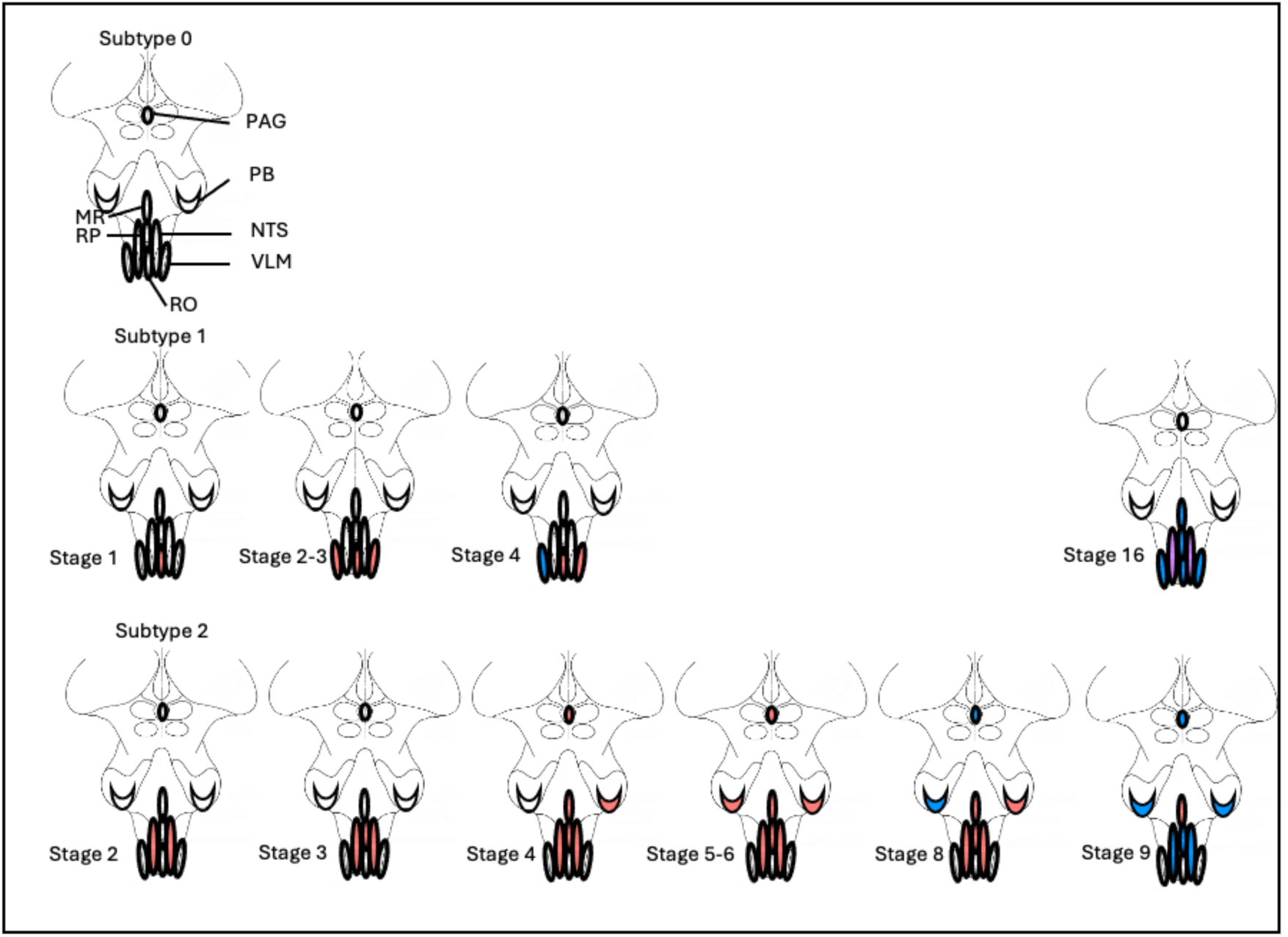
Subtypes and subtype stages of the autonomic control system. Subtype 0 represents normal aging. Subtypes 1 and 2 represent atypical aging. PAG, periaqueductal gray, PB, parabrachial nucleus, MR, raphe magnus, RP, raphe pallidus, RO, raphe obscurus, NTS, nucleus tractus solitarius, VLM, ventrolateral medulla. Red indicates volume losses, i.e., age corrected z-scores >1 but < 2, blue indicates volume losses with age corrected z-scores ≥ 2 but < 3, violet indicates volume losses with age corrected z-scores ≥ 3. Please see main text for a description of the pattern.

The mean systolic blood pressure was not different between the 3 subtypes (subtype 0: mean (SD) 126.4(14.9), subtype 1: mean (SD) 129.5 (16.7), subtype 2: mean (SD) 129 (16.7) mmHg, ANOVA p = 0.094). The same was not true for the mean diastolic blood pressure though (ANOVA p = 0.013). Subtype 1 had a higher diastolic blood pressure (mean (SD): 81.3 (10.2) mmHg) than 2 (mean (SD): 78.1 (10.1) mmHg), p = 0.033 or subtype 0 (mean (SD): 78.2 (10.4) mmHg), p =0.015.

## 4. Discussion

The study has two major findings: 1. Age related volume loss in brainstem structures known to be involved in cognition, motor and autonomic control had generally a negative impact on these functions. Age alone typically explained a larger percentage of the variation of the behavioral variables than the volumes of the brainstem nuclei supporting them and most of the volume-function associations were weaker or lost after accounting for age. The exception were volume losses in a subset of the nuclei (RG, CR) that are involved in motor control where volume loss was associated with increased grip strength. 2. SuStaIn identified 3 to 4 subtypes for each of the four systems. One representing normal aging and 2-3 with distinct patterns representing atypical aging with mild volume losses in the early stages that became more widespread and/or severe in later stages. The mean ages of participants assigned to typical and atypical aging subtypes did not differ and each subtype included participants from the whole HCA age range. Although function in atypical aging subtypes was usually slightly worse than that seen in the typical aging subtype these differences did mostly not reach significance. Considering that HCA focuses on normal aging, i.e., only enrolled subjects who function within the normal range for their age, this was expected.

The implication of these findings will be discussed in more details in the following sections. SuStaIn has been developed and successfully used for the identification of different phenotypes (subtypes) and degrees of their expression (stages) in individuals from heterogeneous patient populations, e.g., patients suffering from different types of fronto-temporal dementia (Young), schizophrenia (Sone et al., 2024) or epilepsy (Lee et al., 2024). Here, SuStaIn was used to characterize volume patterns of brainstem nuclei in a population of normally aging subjects. Because the volumes were corrected for age effects, other factors influencing brain volumes are likely to be responsible for the existence of these additional subtypes. One possibility is that the subtypes represent normal variants of brainstem morphology and the stages different degrees to which a variant is expressed. This interpretation implies that these subtypes have no or only minimal impact on the function supported by the subsystem. The observations that the behavioral variables associated with the subsystems were mostly not significantly different between subtypes and that the volume differences between subtypes were mild ( mean z-scores <2) support this interpretation. Another possibility is that these subtypes represent the earliest manifestations of a neurodegenerative process starting in the brainstem. Some neurodegenerative diseases, e.g., Alzheimer’s (AD) or Parkinson’s disease (PD) (Rueb et al., 2000, Grinberg et al., 2009), can affect the brainstem in a very early, asymptomatic phase or even start there. This means that even a normally aging population could contain individuals in the preclinical stages in whom affected brainstem systems could already differ in subtle ways from those of their non-affected peers. The finding that between 1 – 16% of the subjects assigned to an atypical subtype, some among them still in their thirties or forties, had significantly smaller brainstem volumes (z-scores >2) than those assigned to the typical subtype, supports this interpretation. Finally, these two interpretations are not mutually exclusive, i.e., some subtypes could represent normal and others pathological variants. A longitudinal study is the only way to answer these questions decisively.

SuStaIn identified two subtypes representing atypical aging within the cognitive brainstem system. Subtype 1 was characterized by mild volume loss in MedR, DR and PAG that was already present in the early stages whereas VTA and LC stayed intact until the late stages. PAG, MedR and DR are tightly interconnected (Lovick, 1991). Originally it was thought that they are mostly involved in fear/avoidance processing and association learning (Deakin and Graeff, 1991, Spiacci et al., 2012, Groessl et al., 2018). However, more recent studies demonstrated that their prefrontal projections exert a major influence on executive functions such as cognitive flexibility and patience (Sargin et al., 2019). In subtype 2, mild volume loss started in LC before it spread to the VTA in stage 3. The volume loss became more pronounced in later stages but stayed confined to LC and VTA. Connections between LC and VTA with the hippocampus regulate hippocampal synaptic plasticity and by extension working and episodic memory (Dahl et al., 2023, Duszkiewicz et al., 2019, Nobili et al., 2017, Hagena et al., 2025). DR/MedR and LC/VTA are among the brainstem structures that can accumulate atypical soluble tau very early, in some cases even in early adolescence. At this early state this has no obvious impact on function (Braak et al., 2011) but this changes in later life with increasing Braak stages (Theofilas et al., 2018, Rueb et al., 2000, Grinberg et al., 2009, Lorke et al., 2006). Given this background, it is tempting to speculate that the mild LC/VTA and DR/MedR volume losses observed in this study could represent the earliest stage of a progressive tauopathy, e.g., AD, and that the two subtypes represent different ways how the first signs present themselves. In subjects with a subtype 1 pattern psychiatric symptoms such as depression and anxiety would precede cognitive symptoms (Khan et al., 2023, Khan et al., 2024) and in subjects with a subtype 2 pattern memory and executive problems would dominate the clinical picture.

In addition to the pyramidal tract whose fibers pass through the brainstem to either directly connect with the lower motoneurons in the cranial motor nuclei of the brainstem or the spinal cord, the brainstem contains two other motor control systems. The first is the reticulospinal system that is thought to regulate posture and muscle tone as well as force modulation and locomotion initiation/termination (Brownstone and Chopek, 2018, Danielson et al., 2024, Glover and Baker, 2022, Dautan et al., 2021, Takakusaki et al., 2016, Kim et al., 2017). It consists of the CR and PP that together form the mesencephalic locomotion center (MRC), and the RPO and RPC in the pons and the RG in the medulla (Brownstone and Chopek, 2018). Subtype 2 is consistent with volume loss in the reticulospinal system that starts in its medullary components before spreading to its other parts. The RPO projects onto the RG that controls the activity of the spinal cord motoneurons in a state-dependent manner. In the awake and non-REM sleep state the RPO inhibits the RG and suppresses its inhibitory influence on the motoneurons of the spinal cord. During REM sleep, the MRC suppresses the RPO activity. The disinhibited RG suppresses spinal motoneuron activity which prevents the sleeper from acting out dreamed activities (Xi et al., 2001). It is conceivable that volume losses in RPO and RG disturb this interaction resulting in a suboptimal control over spinal cord motoneurons which could manifest itself in an impaired strength modulation in the awake state and an insufficient movement suppression in REM sleep. Both symptoms can be found in a more pronounced form in early PD (Jo et al. 2015, St Louis et al. 2017).

The second motor control system is the rubro-olivary system that encompasses the OI and NR and is part of the dentato-rubro-olivio loop (Stacho et al., 2024). It is involved in the coordination of limb movements and postural control during walking and facilitates motor learning by relaying incoming signals to the cerebellum (Farhani et al., 2026, Basile et al., 2021). Subtype 1 was characterized by volume loss starting in OI that spread to PP and RPC and Subtype 3 by volume loss in SN, NR and RPO, i.e., both subtypes showed volume loss in core components of the rubro-olivary tract . Damage to the cerebellar and rubral parts of the dentato-rubro-olivary loop typically results in a hypertrophic degeneration of the OI (Wang et al., 2019). This would be consistent with the Subtype 3 pattern. However, without information about potential cerebellar abnormalities a definitive assignment of either of the two subtypes to the dentato-rubro-olivio loop seems premature.

The autonomic brainstem is involved in all aspects of autonomic control, i.e., respiration, thermoregulation, micturition, digestion, cardiovascular function etc. (Schenberg et al, 1994, Guyenet and Stornetta, 2022, Koulu et al., 1986). Blood pressure however was the only autonomic function assessed in the HCA project. It was elevated in the study population as a whole but with the exception of a subgroup whose mean systolic blood pressure was elevated despite antihypertensive treatment not within the hypertensive range. Considering the HCA inclusion criteria, the most likely explanation for the elevated blood pressure is preclinical-clinical essential hypertension. The patho-mechanisms of essential hypertension are not fully understood but likely complex and multifactorial (Carthy, 2013, Coffman, 2011, Jennings et al., 2020). SuStaIn identified two atypical aging patterns. Volume loss in subtype 2 was widespread, i.e., included in the lower brainstem NTS, MR and RP as well as PP in the pons and PAG in the mesencephalon. Volume loss in subtype 1 was constrained to the lower brainstem, i.e., started in the RO in stage 1 and involved VLM in stage 2. The proportion of participants taking antihypertensive medication was with 10-13% about equal in all three subtypes. However, the diastolic blood pressure in subtype 1 was significantly higher than in 0 or 2 and within the hypertensive range. The systolic blood pressure was higher in both atypical aging subtypes compared to the typical aging subtype but not significantly so. Taken together, the findings of this study suggest that volume losses within autonomic brainstem structures could play a role in the development of essential hypertension.

This study has limitations. 1. The HCA project is part of the Human Connectome Lifespan project that also includes the HCP-Development project (ages 5–21 years). The HCP-Aging and HCP-Development use the same imaging protocol. The age range 21–35 years is covered by the HCP Young Adults project that used a slightly different imaging protocol, i.e., the T1 and T2 sequences have a higher resolution (0.7 mm isotropic instead of 0.8 mm isotropic) which complicates combining it with HCP-Aging and HCP-Development particularly when the focus of interest is small structures such as brainstem nuclei. It was therefore not possible to model the structure – age relationship across the whole lifespan. 2. Although HCA participants underwent extensive behavioral testing, its focus was on domains known to be affected by age, in particular cognition and physical fragility. This means that the behavioral data relating to some of the systems of interest for this study was limited, e.g., autonomic function. It is possible that additional information, e.g., heart rate variability, would have provided additional insights regarding function/structure relationship of the different autonomic systems.

Taken together, the findings presented here support the notion that aging affects the brainstem and that changes within the different brainstem sub-systems controlling cognition, motor agility, autonomic function and sleep contribute to the impact aging has on these functions. The spatiotemporal clustering approach implemented in SuStaIn detected different patterns of volume loss or subtypes within each of the investigated brainstem subsystems. Without a longitudinal study it is not possible to decide if these subtypes represent normal variants of brainstem morphology or the first stages of a neurodegenerative process that starts in the brainstem. If the latter, the imaging analysis techniques like the one used in this study could be used to identify individuals at risk for a neurodegenerative disease years before it the first clinical symptoms appear.

## Notes

### Competing Interest Statement

The authors have declared no competing interest.

