## Supplementary figures and images for "Patterns of Typical and Atypical Age-related Brainstem Volume losses"

### Supplemental Tables

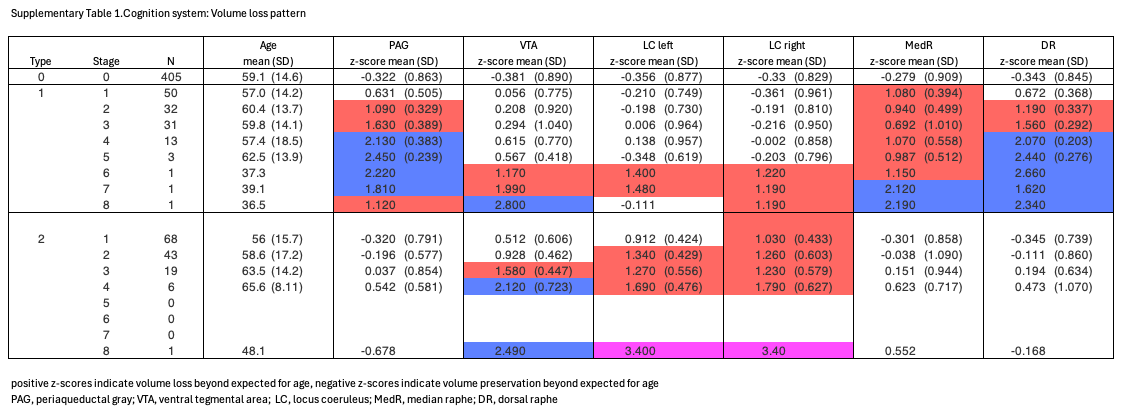

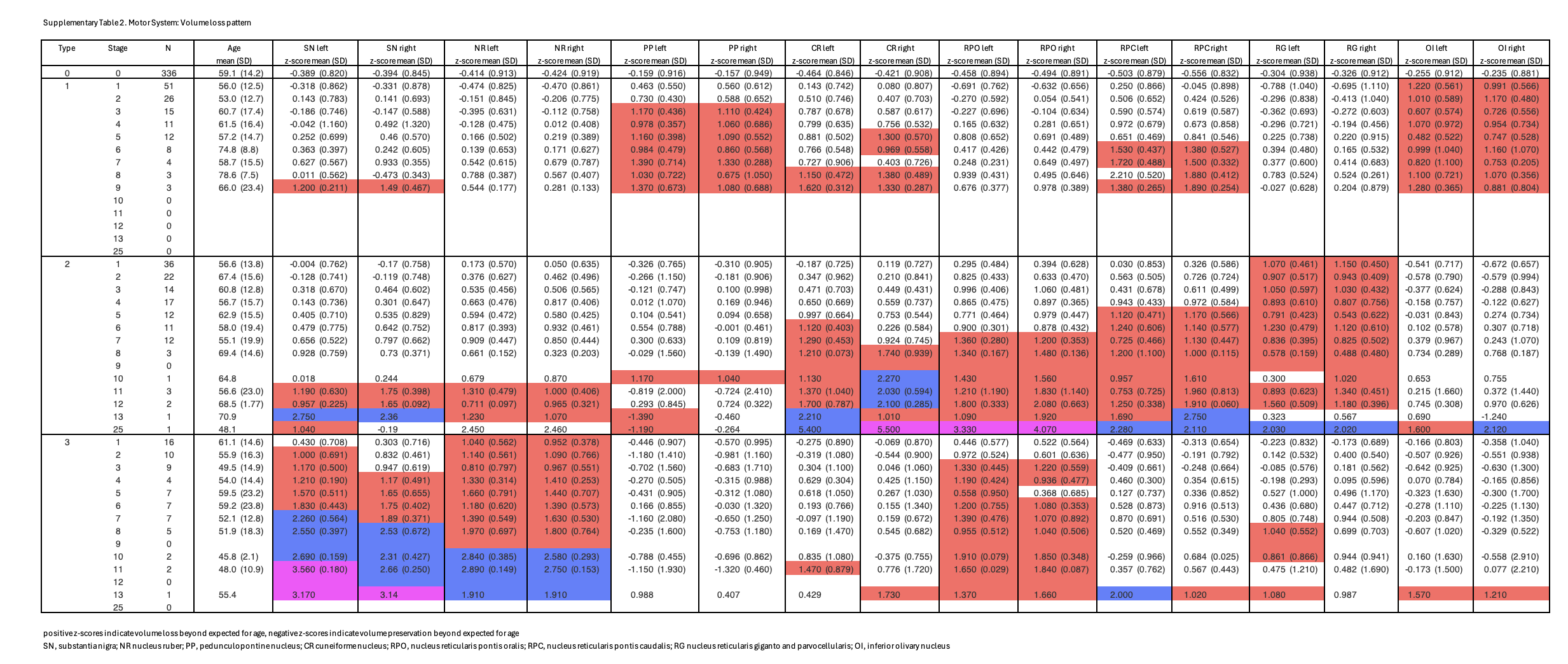

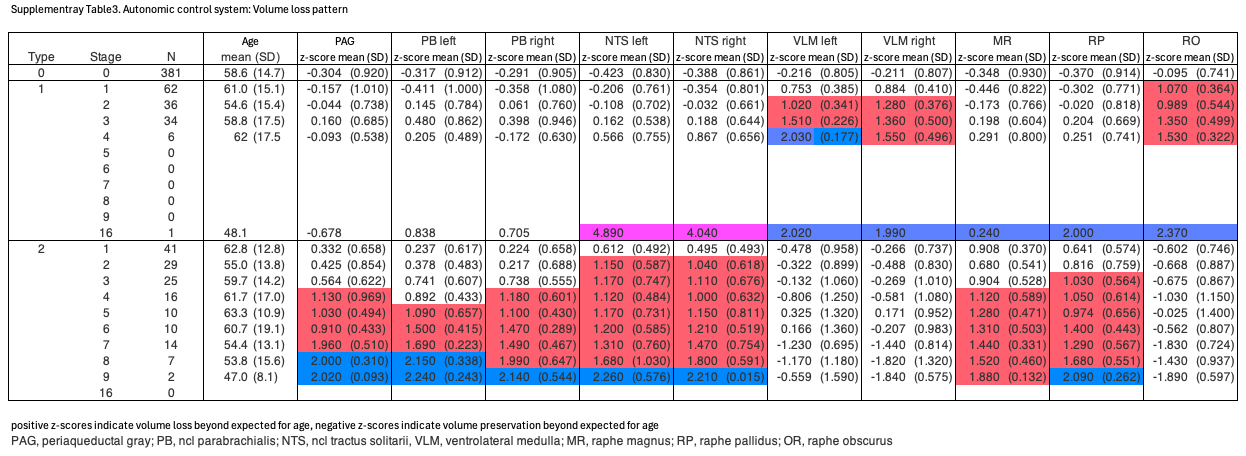
